# MolGene-E: Inverse Molecular Design to Modulate Single Cell Transcriptomics

**DOI:** 10.1101/2025.02.19.638723

**Authors:** Rahul Ohlan, Raswanth Murugan, Li Xie, Vikas Nallabolu, Mohammedsadeq Mottaqi, Shuo Zhang, Lei Xie

## Abstract

Designing drugs that can restore a diseased cell to its healthy state is an emerging approach in systems pharmacology to address medical needs that conventional target-based drug discovery paradigms have failed to meet. Single-cell transcriptomics can comprehensively map the differences between diseased and healthy cellular states, making it a valuable technique for systems pharmacology. However, single-cell omics data is noisy, heterogeneous, scarce, and high-dimensional. As a result, no machine learning methods currently exist to use single-cell omics data to design new drug molecules. We have developed a new deep generative framework named MolGene-E to tackle this challenge. MolGene-E combines two novel models: 1) a cross-modal model that can harmonize and denoise chemical-perturbed bulk and single-cell transcriptomics data, and 2) a contrastive learning-based generative model that can generate new molecules based on the transcriptomics data. MolGene-E consistently outperforms baseline methods in generating high-quality, hit-like molecules on gene expression profiles from two evaluation settings: CRISPR knock-out perturbation profiles from L1000toRNAseq dataset, and single-cell gene expression profiles from Sciplex-3 dataset, both in zero-shot molecule generation setting. This superior performance is demonstrated across diverse *de novo* molecule generation metrics. Extensive evaluations demonstrate that MolGene-E achieves state-of-the-art performance for zero-shot molecular generations. This makes MolGene-E a potentially powerful new tool for drug discovery.

## 1 Introduction

Capitalizing on the success of deep learning across various domains such as natural language, images, and videos, deep generative models have been extensively applied to the generation of small organic compounds targeting a specific disease gene for drug discovery [1]. However, this one-drug-one-target paradigm has had limited success in tackling polygenic, multifactorial diseases. Due to the high costs, prolonged development timelines, and low success rates associated with target-based drug discovery, there has been a resurgence of interest in phenotypic drug discovery. As a matter of fact, approximately 90% of approved drugs have been discovered through a phenotype-driven approach [2]. Therefore, phenotype-based molecular generation, also known as inverse molecule design, holds promise for the discovery of novel therapeutics aimed at addressing medical needs that conventional target-based drug discovery paradigms have failed to meet.

The effectiveness of phenotype-based drug discovery relies upon the careful selection of an appropriate phenotype readout. Chemical-induced transcriptomics has been embraced as a comprehensive, systematic measurement for phenotype drug discovery. The transcriptomic change resulting from chemical exposure can function as a chemical signature for predicting drug responses as well as aid in the elucidation of drug targets and the inference of drug-modulated pathways. This approach has demonstrated successful applications in phenotype drug repurposing [3][4]. Several deep learning methods have been proposed to leverage chemical-induced bulk gene expression data for inverse molecule design. Notably, MolGAN [5] generates molecules conditioning a generative adversarial network with bulk transcriptomics data. Although it shows promising results, we observe that generative adversarial networks (GANs) are susceptible to scalability, as we show that their performance drops significantly when trained on higher dimensional data. Furthermore, GANs have a black-box nature, and inferring the relation between the condition (gene expression) and generation (molecules) is quite cumbersome. Another recent work is the GxVAEs [6], which employs two joint variational autoencoders (VAEs) to facilitate the extraction of latent gene expression features and use it as a condition to generate molecules using a second VAE. However, GxVAEs has not been developed for single-cell transcriptomics data.

The abundance of single-cell omics data provides new opportunities for phenotype-based drug discovery. Single-cell transcriptomics data offer new insights into disease heterogeneity within and across species, illuminating the complexity of pathological processes. An effective therapy often needs to modulate disease etiology at the single-cell level [7]. Furthermore, precise characterization of single-cell chemical transcriptomics is crucial to bridge translational gaps between disease models (e.g., organoids and animals) and human patients, a critical bottleneck in drug discovery [8]. Nonetheless, there remains a scarcity of methods for leveraging single-cell transcriptomics data in inverse molecule design.

Compared with protein structures that exhibit a relatively clean nature, omics data is plagued by its high-dimensionality and susceptibility to noise, stemming from biological stochasticity and technical artifacts. These complexities pose hurdles for single-cell inverse molecule design, exacerbated by the limited availability of chemical-perturbed single-cell transcriptomics data. LINCS1000 [9] serves as a comprehensive chemical transcriptomics database, profiling 19,811 chemicals across 77 cell lines. However, this database profiles only 978 landmark genes. Moreover, the gene expression data in LINCS1000 is obtained using a specific imaging technique, leading to significant distributional discrepancies from RNA-seq data [9]. Due to these challenges, no methods exist for inverse molecule design based on single-cell omics data.

To address these challenges, we introduce MolGene-E, a deep learning framework for single-cell molecule generation. The key contributions of MolGene-E are twofold: First, we develop a domain adaptation model that is capable of harmonizing and denoising L1000toRNAseq [10] and Sciplex-3 [11] single-cell chemical transcriptomics data. Second, we design a generative algorithm that leverages contrastive learning to align phenotypic representations to chemical representations, by integrating these components, MolGene-E facilitates the generation of novel molecules with specific phenotypic traits. Extensive evaluations demonstrate that MolGene-E achieves state-of-the-art performance, positioning it as a potentially powerful new tool for drug discovery.

## 2 Results

To comprehensively evaluate MolGene-E, we design two distinct evaluation scenarios: (i) molecule generation conditioned on target-perturbed gene expression profiles from CRISPR knock-out experiments, and (ii) molecule generation conditioned on chemically induced single-cell gene expression profiles from the SciPlex-3 dataset. Both evaluations are performed in a zero-shot setting, where the model generates molecules for previously unseen gene expression profiles without any fine-tuning, demonstrating its ability to generalize across diverse perturbation types and transcriptomic contexts.

### 2.1 Overview of MolGene-E

MolGene-E is a deep generative framework designed to generate novel drug-like molecules from single-cell transcriptomics data. The pipeline draws inspiration from OpenAI’s DALL-E 2 model [12], which generates images from textual descriptions through a two-step encoding and generative process. Similarly, MolGene-E employs a cross-modal learning approach, where gene expression profiles are aligned with molecular representations to enable de novo molecule generation.

As shown in Figure 1, the framework involves a five-step process that integrates diverse data representations and deep learning modules to align chemical and gene expression profile information effectively.

**Fig. 1:**
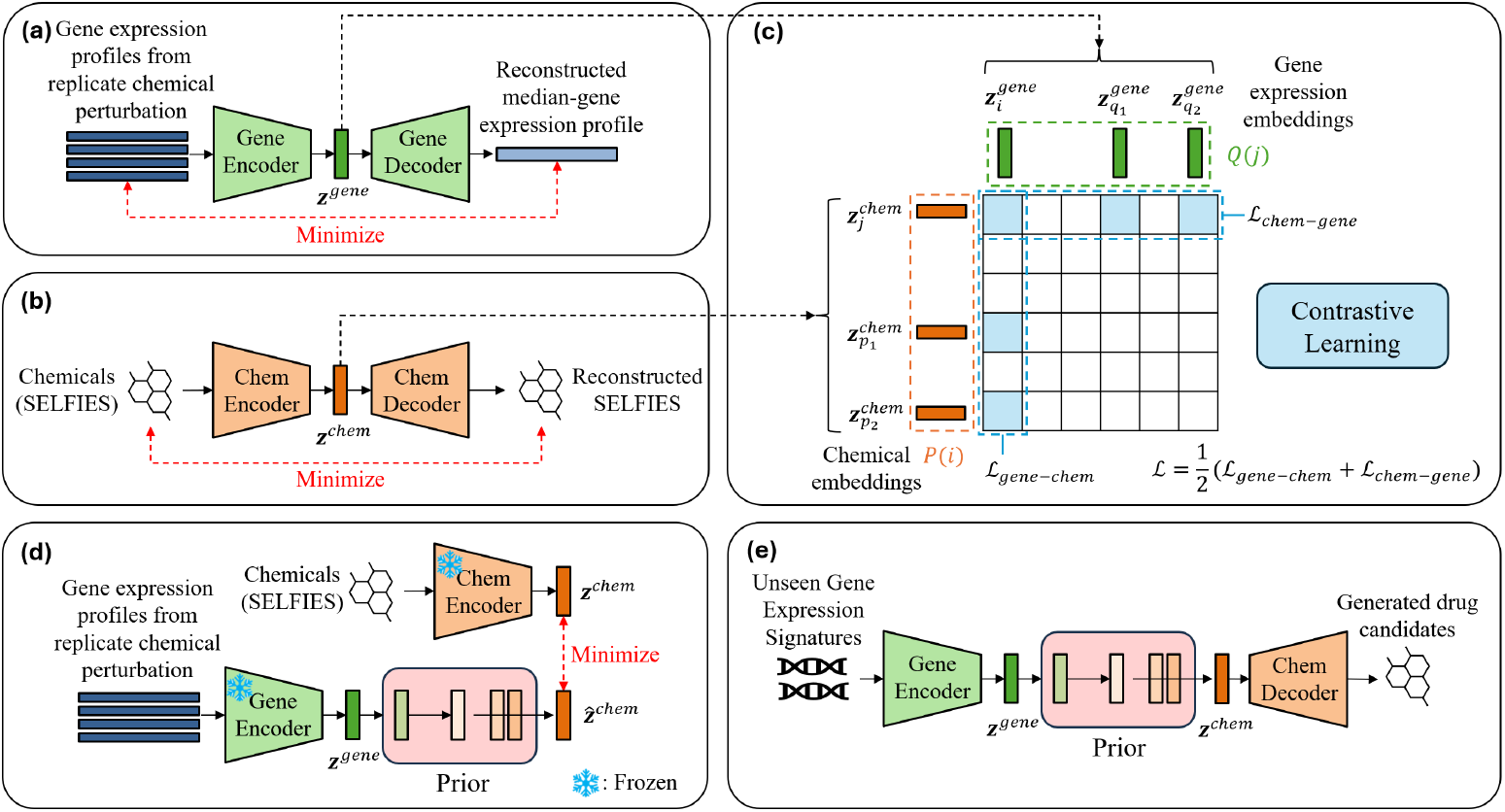
**(a)** A Variational Autoencoder (VAE) denoises the gene expression profiles corresponding to replicate chemical perturbation in a batch by reconstructing their median gene expression profiles. **(b)** We represent chemical structures via SELFIES and use a pretrained frozen VAE to extract the chemical embeddings. **(c)** The gene expression encoder is fine-tuned to align gene embeddings 𝓏^*gene*^ to the chemical embeddings 𝓏^*chem*^ via a contrastive learning module. A supervised objective ℒ (Equation 3) is optimized to maximize the agreement between positive pairs while minimizing the similarity between negative pairs. **(d)** A prior model is trained to map the inferred 𝓏^*gene*^ to the inferred 𝓏^*chem*^. **(e)** Given unseen gene expression profiles, the inferred 𝓏^*gene*^ are mapped to 𝓏^*chem*^ via the pretrained prior model, and are further decoded using the SELFIES VAE’s decoder to generate drug candidates.

First, a Variational Autoencoder (VAE) denoises gene expression profiles corresponding to replicate chemical perturbations by reconstructing their median gene expression profiles (Figure 1a). This step ensures robustness against biological noise in transcriptomic data. Next, chemical structures are represented using SELFIES and passed through a pretrained, frozen VAE to obtain chemically meaningful embeddings (Figure 1b).

A contrastive learning module is then employed to align the gene expression space with the chemical space (Figure 1c). The gene expression encoder is fine-tuned to maximize the agreement between gene embeddings 𝓏^*gene*^ and their corresponding chemical embeddings 𝓏^*chem*^ while minimizing similarity with negative samples. This supervised contrastive loss optimizes the model’s ability to infer chemical perturbations from gene expression data. Positive pairs are constructed by pairing each gene expression profile 𝓏^*gene*^ with its corresponding chemical embedding 𝓏^*chem*^ derived from the molecule used in the perturbation experiment. To account for chemical replicates where multiple gene expression profiles correspond to the same perturbation molecule, all such profiles are treated as additional positives for a given anchor. Negative pairs are implicitly formed by all other gene-molecule pairs within the training batch. The contrastive loss is computed symmetrically in both directions (gene →molecule and molecule → gene) and reduces to the standard InfoNCE loss [13] in the absence of replicates.

To enable molecule generation, a prior model is trained to learn the mapping from 𝓏^*gene*^ to 𝓏^*chem*^ (Figure 1d). Given an unseen gene expression profile and the encoded gene embedding 𝓏^*gene*^, the trained prior model predicts its corresponding 𝓏^*chem*^, which is subsequently decoded by the SELFIES VAE to generate novel molecular structures (Figure 1e).

By harmonizing multi-modal data and leveraging generative modeling principles, MolGene-E provides a powerful approach for discovering drug candidates based on transcriptomic responses, advancing the field of AI-driven drug discovery.

### 2.2 MolGene-E Improves the Success Rate of Inverse Molecule Design

Figure 2 compares the reconstructed gene expression values from the denoising VAE (Figure 1a) in MolGene-E with the median gene expression values derived from replicate samples of a perturbation in the validation and test set of L1000toRNAseq dataset. The histogram shows the density distribution of gene expression values, where the blue bars represent the original median gene expression values, and the orange bars represent the reconstructed values. The close overlap of the two distributions on both the validation and test sets indicates that the denoising VAE in MolGene-E effectively captures and reconstructs the original distribution of gene expression values, even in the presence of noise from biological or experimental variability.

**Fig. 2:**
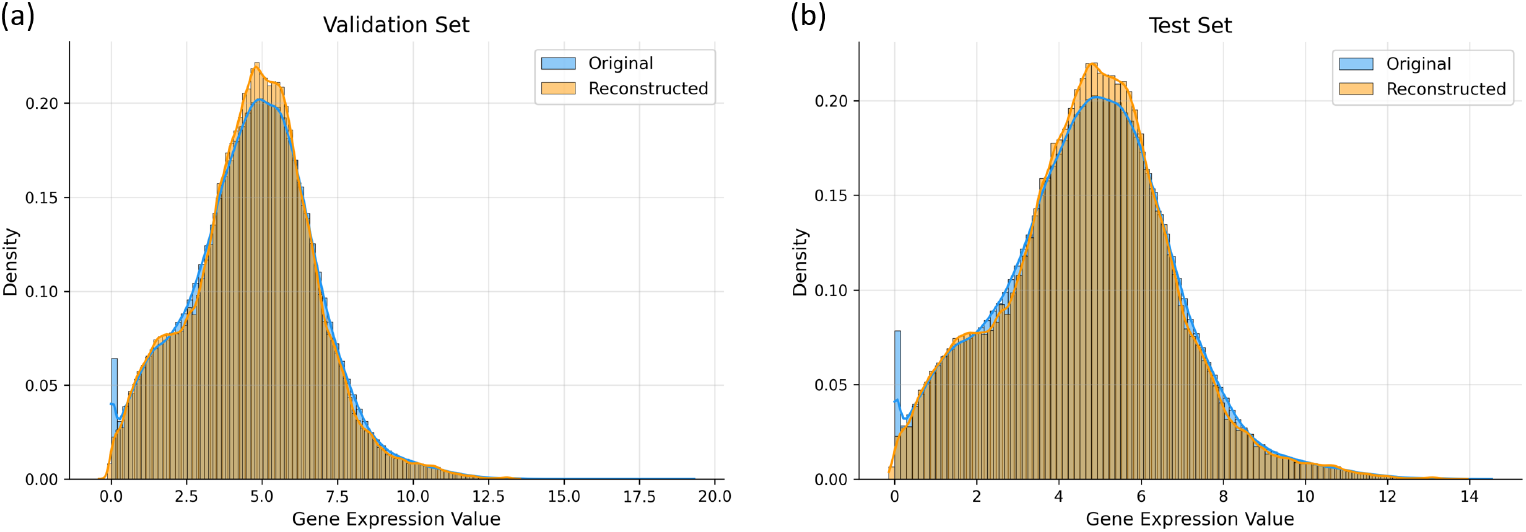
Reconstruction performance of the denoising VAE on the **(a)** validation set and **(b)** test set of L1000toRNAseq dataset.

Then we use a challenge task to evaluate the performance of the molecular generation from gene expressions. If a drug can correctly revert gene expressions from a disease state to a healthy state, the drug could interact with disease-causing genes, i.e., drug targets. In other words, the gene expression changes that are caused by the target gene knock-out or knock-down should be similar to those that result from the chemical perturbation targeting the knock-out/down gene. In our experiments, reference molecules were drawn from the test-split of L1000toRNAseq dataset, restricted to compounds with single-gene targets. For each target gene, we identified the corresponding CRISPR knock-out expression profile from the same dataset in the MCF7 cell line [14]. These knock-out gene expression profiles were then used as conditioning inputs for inference of drug candidates, and the generated molecules were compared against the reference compounds targeting the same gene.

To evaluate the quality of generated molecules, we first consider basic metrics, including **Validity, Uniqueness, Novelty**, and **Diversity**, as listed in Table 1. MolGene-E demonstrates superior performance across most basic metrics compared to the baseline models, MolGAN and GxVAE. It achieves perfect scores (1.00) for validity, uniqueness, and novelty, highlighting its robust capability to generate valid, unique, and novel molecular candidates. For diversity, MolGene-E achieves a score of 0.91, matching GxVAEs and outperforming MolGAN (0.88), indicating that it maintains a high level of structural variability among generated molecules.

**Table 1:** Evaluation metrics of molecules generated from CRISPR knock-out gene expression profiles. We mark the best results in bold and the second-best results with underline.

| Model | Validity ↑ | Uniqueness ↑ | Novelty ↑ | Diversity ↑ | QED ↑ | SA ↓ |
| --- | --- | --- | --- | --- | --- | --- |
| MolGAN | <u>0.29</u> | <u>0.99</u> | **1.00** | <u>0.88</u> | <u>0.68</u> | 4.06 |
| GxVAEs | 0.27 | 0.96 | <u>0.95</u> | **0.91** | 0.59 | **3.00** |
| MolGene-E | **1.00** | **1.00** | **1.00** | **0.91** | **0.69** | <u>3.59</u> |

Since chemically valid and diverse molecules may not necessarily have favorable drug-related properties, we further evaluate drug-likeness (**QED**) and synthesizability (**SA**), as listed in Table 1. For QED, MolGene-E achieves the highest score of 0.69, surpassing MolGAN (0.68) and GxVAEs (0.59), indicating that MolGene-E generates more drug-like molecules. While MolGene-E’s SA score of 3.54 is higher (worse) than GxVAEs (3.00) but lower (better) than MolGAN (4.06), it remains within a reasonable range and is accompanied by strong performance across the basic metrics and superior drug-likeness. Overall, MolGene-E demonstrates strong and consistent performance in generating chemically valid, unique, novel, and drug-like molecular candidates from CRISPR knock-out gene expression profiles.

Beyond evaluating basic generative metrics and drug-related properties, we further assess how well the generated molecules preserve structural similarity to reference compounds. As shown in Figure 3, we compare MolGene-E with the baseline models, MolGAN and GxVAEs, in terms of similarity distributions between generated and reference molecules. We employ multiple similarity measures, including Tanimoto similarities based on MACCS keys [15], Fraggle similairty [5], and cosine similarities based on Reduced Graphs (ErG) fingerprints [16], to capture complementary aspects of molecular similarity, such as substructure patterns, fragment-level similarity, and pharmacophore-related topology. Across all three similarity measures, MolGene-E (green) consistently achieves higher mean similarity compared to MolGAN and GxVAEs, as evidenced by the rightward shift of its distribution and the position of its mean (dashed line), with statistically significant improvements (one-sided Mann–Whitney U-test, p-value *<* 0.05; see Appendix C Table C1 for details). This consistent superiority across diverse fingerprint schemes demonstrates that MolGene-E generates molecules that are structurally more similar to reference compounds regardless of the molecular representation used, highlighting the robustness of our approach.

**Fig. 3:**
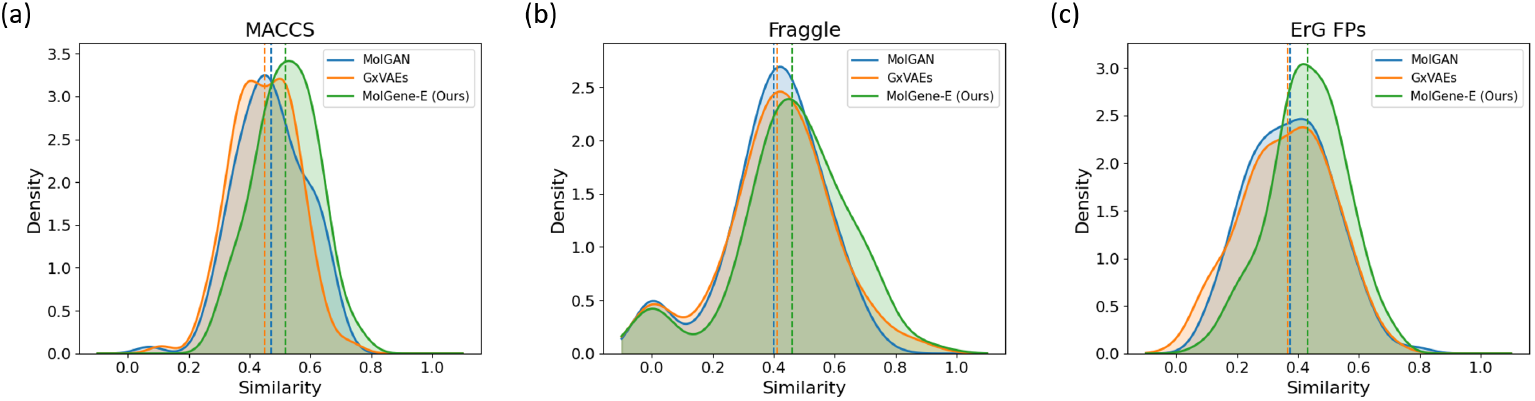
Comparison of similarity distributions between hit-like molecules generated from CRISPR knock-out gene expression profiles and reference molecules. **(a)** Distributions of Tanimoto similarities based on MACCS keys. **(b)** Distributions of Fraggle similarities. **(c)** Distributions of cosine similarities based on ErG fingerprints.

Figure 4 presents representative examples of reference and generated molecule pairs from CRISPR knock-out gene expression profiles, with the top four pairs selected under each of three similarity measures: MACCS key Tanimoto similarity, Fraggle similarity, and ErG fingerprint cosine similarity. The top-scoring pairs under each metric demonstrate that MolGene-E generates molecules with meaningful structural resemblance to the reference compounds. In addition, the generated molecules exhibit reasonable drug-like properties, with many examples showing relatively high QED values (often above 0.7) and low synthetic accessibility scores (typically below 3), indicating favorable drug-likeness and synthesizability. These results further validate its ability to produce pharmacologically relevant candidates conditioned on gene expression perturbations.

**Fig. 4:**
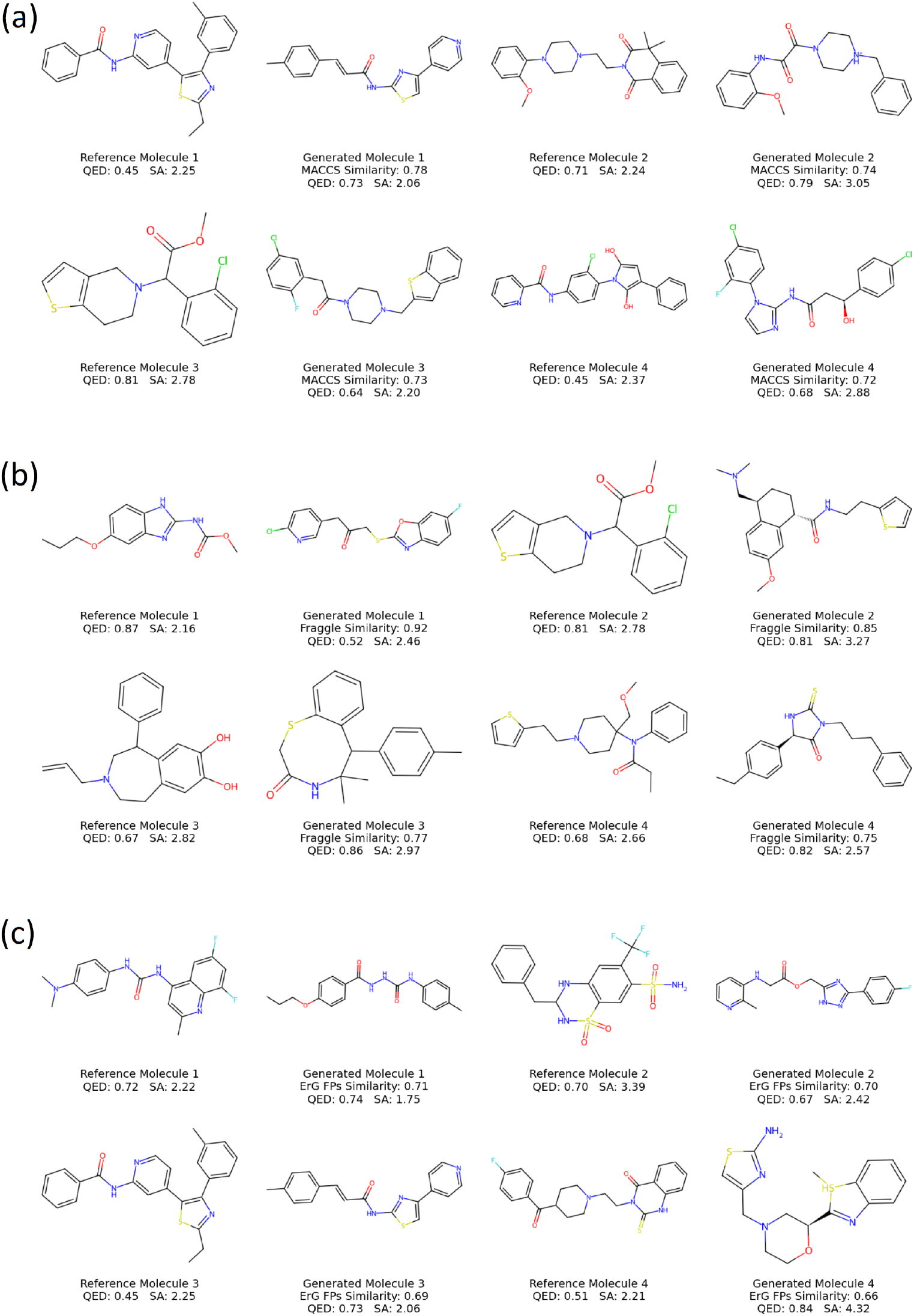
Examples of reference and generated molecules from CRISPR knock-out gene expression profiles. Top pairs based on **(a)** Tanimoto similarity using MACCS keys, **(b)** Fraggle similarity, and **(c)** cosine similarity using ErG fingerprints.

### 2.3 MolGene-E Can Be Applied to Single-Cell Data

While MolGene-E is trained on bulk transcriptomics data from the L1000toRNAseq dataset [10], direct training on single-cell perturbation data is infeasible due to the limited scale of available datasets such as SciPlex-3 [11], which contains only 188 compounds. We therefore evaluate MolGene-E in a zero-shot setting on single-cell data to assess its ability to generalize across transcriptomic modalities. To bridge the representation gap between bulk and single-cell transcriptomics, SciPlex-3 profiles are harmonized into the L1000toRNAseq representation space prior to inference, enabling direct application of the model without retraining. Since each chemical perturbation includes multiple replicates, gene expression profiles are randomly sampled for each perturbation.

The results in Table 2 demonstrate that MolGene-E outperforms MolGAN and GxVAEs across most basic evaluation metrics. For validity, MolGene-E achieves a perfect score of 1.00, substantially higher than MolGAN (0.29) and GxVAEs (0.30), largely due to the use of SELFIES instead of SMILES, which ensures syntactic validity of generated molecules. MolGene-E also attains perfect scores of 1.00 for both uniqueness and novelty. For diversity, MolGene-E achieves a score of 0.91, matching GxVAEs and outperforming MolGAN (0.87), indicating that it maintains high structural variability among generated molecules.

**Table 2:** Evaluation metrics of molecules generated from Sciplex-3 single-cell gene expression profiles. We mark the best results in bold and the second-best results with underline.

| Model | Validity ↑ | Uniqueness ↑ | Novelty ↑ | Diversity ↑ | QED ↑ | SA ↓ |
| --- | --- | --- | --- | --- | --- | --- |
| MolGAN | 0.29 | 0.95 | **1.00** | <u>0.87</u> | <u>0.60</u> | 3.92 |
| GxVAEs | <u>0.30</u> | <u>0.96</u> | <u>0.95</u> | **0.91** | 0.59 | **3.03** |
| MolGene-E | **1.00** | **1.00** | **1.00** | **0.91** | **0.68** | <u>3.62</u> |

In terms of drug-related properties, MolGene-E achieves the highest QED score of 0.68, surpassing MolGAN (0.60) and GxVAEs (0.59), suggesting improved drug-likeness of the generated molecules. While MolGene-E’s SA score of 3.62 is worse than GxVAEs (3.03) but better than MolGAN (3.92), it remains within a reasonable range, reflecting an acceptable trade-off between synthesizability and other performance gains.

Beyond these metrics, we further evaluate how well the generated molecules preserve structural similarity to reference compounds. Figure 5 shows the distributions of similarity scores between generated and reference molecules. MolGene-E consistently achieves higher similarity scores compared to both baseline models across all three similarity schemes (MACCS key Tanimoto similarity, Fraggle similarity, and ErG fingerprint cosine similarity), indicating better structural alignment with reference compounds, with statistically significant improvements (one-sided Mann–Whitney U test, p-value *<* 0.05; Appendix C Table C2).

**Fig. 5:**
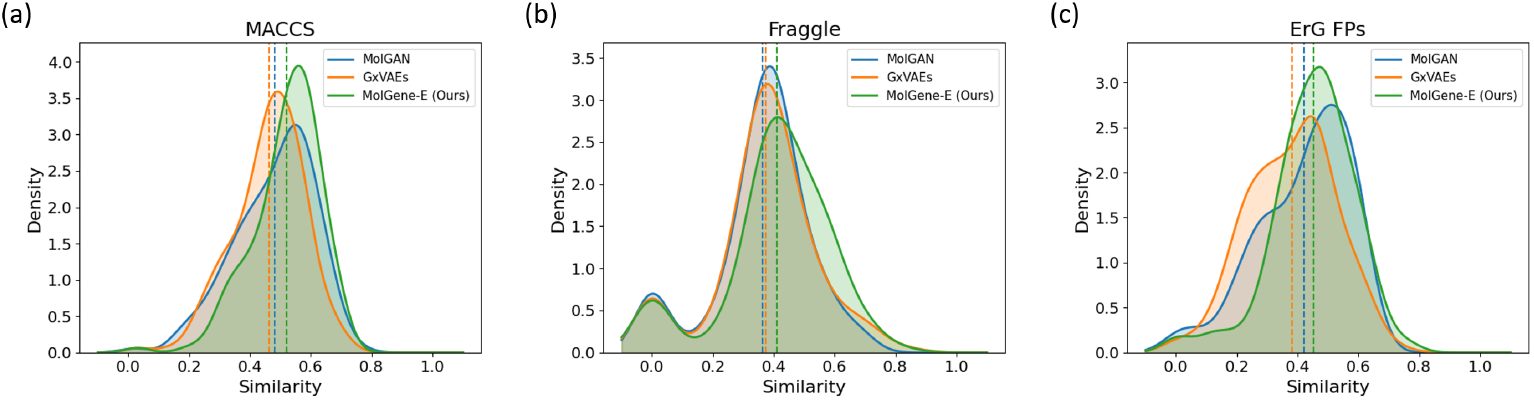
Comparison of similarity distributions between hit-like molecules generated from Sciplex-3 single-cell gene expression profiles and reference molecules. **(a)** Distributions of Tanimoto similarities based on MACCS keys. **(b)** Distributions of Fraggle similarities. **(c)** Distributions of cosine similarities based on ErG fingerprints.

Overall, MolGene-E demonstrates strong and consistent performance in generating chemically valid, diverse, and drug-like molecular candidates from single-cell transcriptomics data. Notably, despite being trained solely on bulk transcriptomics data, MolGene-E generalizes effectively to single-cell settings in a zero-shot manner, highlighting its potential for phenotype-based inverse molecular design across different transcriptomic modalities under limited-data conditions.

### 2.4 Ablation Studies

To evaluate the efficacy of various design choices in our molecular generation pipeline, we conducted comprehensive ablation studies on the Sciplex-3 dataset in a zero-shot setting. The results are discussed below.

#### Effect of the prior model

We remove the prior model in MolGene-E to build an ablated model called MolGene-E_*NP*_ . When the prior model is utilized, gene expression embeddings are first mapped to the latent space, which is then used for molecule generation. Without the prior model, gene expression embeddings are directly used from the CLIP encoder. The molecule generation metrics for this comparison are shown in Table 3. Removing the prior model leads to substantial degradation in performance, including a much worse SA score (5.91 vs 3.62), a markedly lower QED score (0.25 vs 0.68), and reduced structural similarity to reference molecules across both MACCS key Tanimoto similarity (0.47 vs 0.52) and Fraggle similarity (0.39 vs 0.41). These results indicate that the prior model plays a crucial role in aligning gene expression representations with the molecular latent space, thereby enabling the generation of synthetically accessible and drug-like molecules with meaningful structural resemblance to reference compounds.

**Table 3:** Results of ablation study on Sciplex-3 dataset.

| Model | Prior | Pretrain | QED ↑ | SA ↓ | MACCS ↑ | Fraggle ↑ |
| --- | --- | --- | --- | --- | --- | --- |
| MolGene-E<sub>NP</sub> | × | ✓ | 0.25 | 5.91 | 0.47 | 0.39 |
| MolGene-E<sub>E</sub> | ✓ | × | 0.54 | 4.30 | **0.52** | 0.40 |
| MolGene-E | ✓ | ✓ | **0.68** | **3.62** | **0.52** | **0.41** |

#### Effect of pretraining the gene expression encoder for denoising

We further compare MolGene-E with a variant MolGene-E_*E*_, where the gene expression decoder in the denoising VAE is removed, and the gene expression encoder is trained end-to-end with the remaining modules without denoising pretraining. The molecule generation metrics for this comparison are listed in Table 3. Pretraining the encoder leads to improved chemical quality, with a better SA score (3.62 vs 4.30) and higher QED (0.68 vs 0.54), while maintaining comparable structural similarity to reference molecules in terms of MACCS key Tanimoto similarity (0.52 vs 0.52) and Fraggle similarity (0.41 vs 0.40). This suggests that denoising pretraining enhances the robustness of gene expression representations, leading to more reliable molecule generation without compromising structural fidelity.

The results of the ablation studies highlight the importance of both components. The prior model facilitates effective cross-modal alignment between gene expression and molecular representations, while the pretrained encoder improves the quality of generated molecules by providing more robust and noise-aware embeddings. These findings provide insights into the key design choices that contribute to high-quality molecule generation in our framework.

## 3 Discussion

In this paper, we developed a deep generative model that utilizes phenotypic properties from transcriptomic profiles, including single-cell omics data, to generate high-quality lead candidates for drug discovery. MolGene-E consistently outperforms baseline methods in generating high-quality, hit-like molecules from gene expression profiles obtained from CRISPR-based gene knock-out perturbations as well as single-cell datasets. This superior performance is demonstrated across *de novo* molecule generation metrics, including novelty, diversity, uniqueness, drug-likeness, and synthesizability.

MolGene-E demonstrates strong performance across structural similarity metrics, consistently achieving higher similarity between generated molecules and their corresponding reference compounds across multiple complementary measures, including MACCS key Tanimoto similarity, Fraggle similarity, and ErG fingerprint cosine similarity, which together capture substructure patterns, fragment-level similarity, and pharmacophore-related topology. The consistent improvements across these representations indicate that MolGene-E generates molecules that are not only structurally aligned with reference compounds but also more likely to be functionally analogous. This comprehensive multi-perspective evaluation further supports the effectiveness of MolGene-E in generating biologically relevant molecular candidates.

Future work includes incorporating multiple cell lines and conditioning molecule generation on multi-omics data, to better capture the heterogeneity of cellular contexts and enable a more robust framework that reflects complex biological environments *in vivo*. Additionally, expanding the model to integrate diverse datasets will enhance its ability to generalize across different biological contexts, thereby improving its predictive power and utility in identifying effective therapeutic compounds. Such extensions have the potential to enable more precise and context-aware drug discovery, ultimately accelerating the development of new treatments and improving patient outcomes.

## 4 Methods

### 4.1 Denoising VAE for Gene Expression Profiles

In chemical-induced bulk gene expression data, multiple distinct gene expression profiles perturbed by replicate chemicals can exist. In order to manage and interpret the complex data from multiple replicates, MolGene-E employs a VAE for denoising, which is trained with the objective of reconstructing the median gene expression profile from the gene expression profiles corresponding to replicate chemical perturbations in a batch (Figure 1a). The training objective incorporates a standard reconstruction loss and KL divergence loss [17], with dynamic weighting between these components. During training, the weight of the reconstruction component is gradually reduced while the KL divergence component is increased, ensuring that the latent space becomes well-regularized and captures the underlying structure of the data. This approach ensures that the VAE captures the most representative gene expression profile, reducing noise and focusing on the core response to chemical perturbations. This process enhances the reliability of the gene expression data used in further steps.

### 4.2 SELFIES VAE for Chemicals

For representing the chemical structures in perturbations, MolGene-E leverages SELFIES (Self-Referencing Embedded Strings) [18] due to its guaranteed 100% validity in contrast to using SMILES strings to represent molecules for molecule design [5]. These SELFIES strings are encoded using a VAE model pretrained on ZINC dataset [19] (Figure 1b). The use of SELFIES allows for a comprehensive and error-resistant encoding of molecular structures, facilitating seamless integration with machine learning models.

### 4.3 Alignment of Gene Expressions and Chemical Representations

The key innovation in MolGene-E lies in aligning the gene expression profiles with their corresponding chemical perturbations. This is achieved through a contrastive learning module (Figure 1c) trained with a supervised contrastive loss ℒ (Equation 3) inspired by CLIP [20] and SupCon loss [21]. The objective of this module is to align the embeddings of phenotypes (gene expression profiles) with the embeddings of the SELFIES representations of the chemicals that caused the perturbations. It is also specifically designed to deal with the existence of multiple distinct gene expression profiles perturbed by replicate chemicals in a batch. By doing so, MolGene-E ensures that the biological effects of chemicals are accurately reflected in their encoded representations.

### 4.4 Mapping Gene Expressions to Chemical Embeddings

To complete the alignment process, MolGene-E employs a Multi-Layer Perceptron (MLP)-based prior model (Figure 1d). This model is trained to map the embeddings of gene expression profiles to the embeddings of their corresponding chemical counterparts. The MLP-based prior effectively bridges the gap between biological responses and chemical structures, enabling the generation of novel molecules that can induce desired gene expression changes.

### 4.5 Generation of Drug Candidates

After training the prior model, gene expressions corresponding to chemical perturbations can be used for inference to generate drug candidates that might result in similar perturbation effects (Figure 1e). The gene expression embeddings 𝓏^*gene*^ are extracted using the pretrained gene expression encoder in the contrastive learning module and subsequently input to the prior model to compute chemical embeddings 𝓏^*chem*^. 𝓏^*chem*^ are then decoded via the SELFIES VAE model to obtain potential drug candidates in the form of novel molecular structures.

### 4.6 Implementation Details

#### 4.6.1 Datasets

The L1000toRNAseq dataset, originally containing 978 landmark genes, was transformed to RNA-seq-like profiles encompassing 23,614 genes using a cycleGAN model as described by [10]. The dataset includes gene expression profiles from 221 human cell lines treated with over 30,000 chemical and genetic perturbations, resulting in over 3 million expression profiles. We filtered the data for chemical perturbations with 24-hour treatment times and 10 µM dosage for the MCF7 cell line, resulting in 3116 genes with high variance (variance *>* 0.75). For training MolGene-E we did a 80-20-20 split to get training, validation, and test sets while ensuring there was no chemical overlap in the data splits. This dataset was used to train all components of MolGene-E. Specifically, the denoising GeneVAE was trained on the chemical perturbation profiles from the MCF7 cell line to learn to reconstruct median gene expression profiles from replicate measurements. The contrastive learning module (CLIP) was subsequently trained on the same dataset to align gene expression embeddings with their corresponding chemical embeddings derived from the SELFIES VAE. The SELFIES VAE, pretrained on the MOSES dataset [22] and available from the mol opt repository [19], was kept frozen throughout and used to derive chemical embeddings for all training stages. Finally, the MLP prior model was trained on the aligned embeddings from the CLIP module using the same L1000toRNAseq training split. Zero-shot inference was performed on two held-out datasets: CRISPR knock-out perturbation profiles and Sciplex-3 single-cell profiles, neither of which was used during any training stage. Both baseline models MolGAN and GxVAEs were retrained from scratch using their original architectures on the same MCF7 cell line data splits used for MolGene-E, ensuring a fair and consistent comparison.

The CRISPR Perturbations L1000 dataset, sourced from the sigcom portal [14], consists of 1218 L1000 signatures for 44 different transcription factors (TFs) targeted by CRISPR knock-out perturbations. We used this data for zero-shot inference filtering it for the MCF7 cell line and the samples consisting of single targets in order to avoid confounding effects that may arise from multi-target perturbations, allowing for a clearer evaluation of the model’s ability to infer the effects of individual transcription factor knock-outs.

The Sciplex-3 dataset, sourced from [11] and harmonized by scPerturb [23], includes single-cell transcriptomic profiles of 188 compounds across three cancer cell lines. We focused on the MCF7 cell line to be used for inference since MolGene-E was trained on MCF7 cell line data and filtered the data to improve quality. Since all experiments are performed on a single cell line (MCF7), control group variations are consistent across all samples, making perturbed-minus-control normalization redundant in this setting. Instead, gene expression profiles are z-score normalized using the mean and standard deviation computed from the MCF7 training set, which effectively accounts for baseline expression differences across perturbations. The dataset was harmonized with the L1000toRNAseq data using two steps: first, missing values were imputed using the Deep Count Autoencoder (DCA) [24]; second, a VAE-based domain adaptation model was trained on chemicals overlapping between the Sciplex-3 and L1000toRNAseq datasets, learning to reconstruct the corresponding L1000toRNAseq gene expression profiles from the gene expression profiles measured in the Sciplex-3 dataset. This trained model was then applied to all gene expression profiles in Sciplex-3 to map them into the L1000toRNAseq representation space, as shown in Appendix B Figure B1.

#### 4.6.2 Evaluation Metrics

For performance evaluation, the following measures were used. Structural similarity between reference and generated molecules is evaluated using three complementary molecular representations, including **Tanimoto similarity based on MACCS keys** [15], **Fraggle similarity** [5], and **cosine similarity based on Reduced Graph (ErG) fingerprints** [16], capturing substructure patterns, fragment-level similarity, and pharmacophore-related topology, respectively. Higher similarity scores indicate that the generated molecules are more structurally aligned with their corresponding reference compounds and are therefore more likely to be potential drug candidates.

**Novelty** is the fraction of generated molecules not observed in the training set. **Uniqueness** is the fraction of distinct molecules among all valid generated molecules across all targets. **Diversity** measures the chemical variability among the generated molecules, computed as one minus the average pairwise Tanimoto similarity using Morgan fingerprints (radius=3, 2048 bits) between all generated molecules, with higher diversity indicating a broader exploration of the chemical space. **Validity** and **SA** (synthetic accessibility scores) are computed using the RDKit library [25]. **QED** (Quantitative Estimate of Drug-likeness) [26] measures the drug-likeness of generated molecules, with higher scores indicating more drug-like properties.

#### 4.6.3 Model Settings

For the denoising VAE, we used hidden layers of sizes [512, 256, 128] with layer normalization [27] and a dropout rate of 0.2. We used a latent dimension of 64. The weight for KL-term of loss was increased linearly from the first to the last epoch. We trained the model for 500 epochs.

For the SELFIES VAE, it maximizes a lower bound of the likelihood (evidence lower bound (ELBO)) instead of estimating the likelihood directly. We used the pretrained model from open source molecular optimization benchmark [19].

For the prior model, it is an MLP with hidden sizes [512, 256, 128] and a latent dimension of size 128. The model was trained to minimize a mean squared error loss for the reconstruction of chemical embedding space utilizing the aligned spaces from both modalities. The model used a learning rate of 1e-3 and a batch size of 128.

For the training of MolGene-E, we minimize the contrastive learning objective (Equation 3) with an approach similar to [20]. MolGene-E was trained for 290 epochs with a batch size of 128 and a learning rate of 1e-4. A projection network with MLP hidden layers [128, 128] was further added to the gene expression encoder. When generating drug candidates, 20 unique chemical candidates are generated by sampling from the latent space for each gene expression profile. The molecule with the highest similarity score is selected as the top candidate. For further details on the training (Algorithm 1,2) and inference (Algorithm 3) process, please refer to Appendix A.

To define the contrastive loss, we introduce ℒ_gene-chem_ and ℒ_chem-gene_. The former aligns each arbitrary anchor gene expression embedding 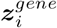 with an index *i* to all corresponding replicate chemical perturbation embeddings 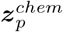 with indices *p* ∈ *P* (*i*) in a batch:

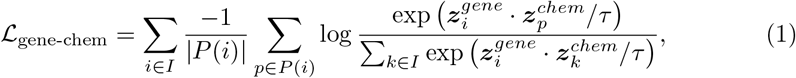

while the latter aligns each arbitrary anchor chemical embedding 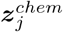 with an index *j* to all corresponding perturbed gene expression embeddings 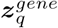 with indices *q* ∈ *Q*(*j*) in a batch:

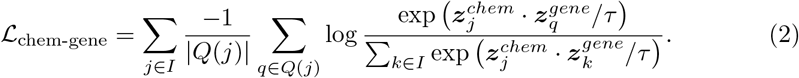

In the two equations above, *τ* denotes the temperature parameter controlling the sharpness of the similarity scores, | *P* | denotes the cardinality of *P*, and *I* is the set of all indices in the batch.

The final contrastive loss ℒ is obtained by combining the two losses above:

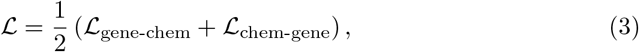

The SELFIES chemical embeddings are used directly from the pre-trained chemical encoder model underscored in the previous section and its parameters are frozen while training.

## Data Availability

The datasets analyzed in the study are publicly available. The L1000RNAtoseq chemical perturbations and CRISPR perturbations can be downloaded from https://lincs-dcic.s3.amazonaws.com/LINCS-data-2020/RNA-seq/cp_predicted_RNAseq_profiles.gctx and https://lincs-dcic.s3.amazonaws.com/LINCS-data-2020/RNA-seq/xpr_predicted_RNAseq_profiles.gctx respectively. The sciplex-3 dataset can be obtained from the scPerturb portal at https://zenodo.org/records/7041849/files/SrivatsanTrapnell2020_sciplex3.h5ad?download=1.

## Code Availability

The code used in this study will be made accessible to reviewers via Code Ocean during the review phase. The scripts and necessary data to reproduce the key results presented in this manuscript will be included.

## Acknowledgement

This project has been funded with federal funds from the National Institute of General Medical Sciences of the National Institute of Health (R01GM122845), the National Institute on Aging of the National Institute of Health (R01AG057555, R21AG083302), and the National Science Foundation (NSF2230354).

## Author Contributions

Lei X. conceived the presented idea and acquired funding. R. O. and R. M. performed the computations, baseline experiments, and verified the analytical methods. M. M. organized the datasets used in the study. Li X. and V. N. conducted the experiments. Lei X. and S. Z. supervised the design of methods and experiments, and the writing of the manuscript.

## Appendix

A Algorithms for Training and Inference

### Algorithm 1

Training the Contrastive Learning Module

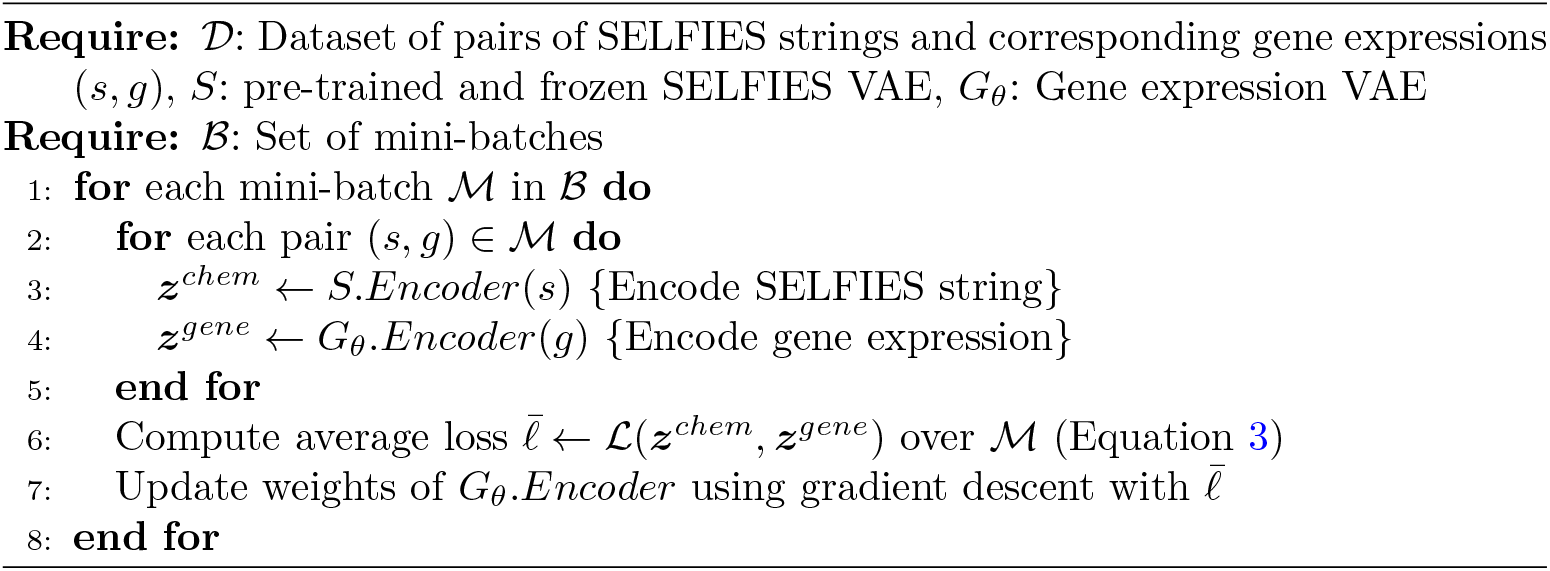

### Algorithm 2

Training the Prior Module

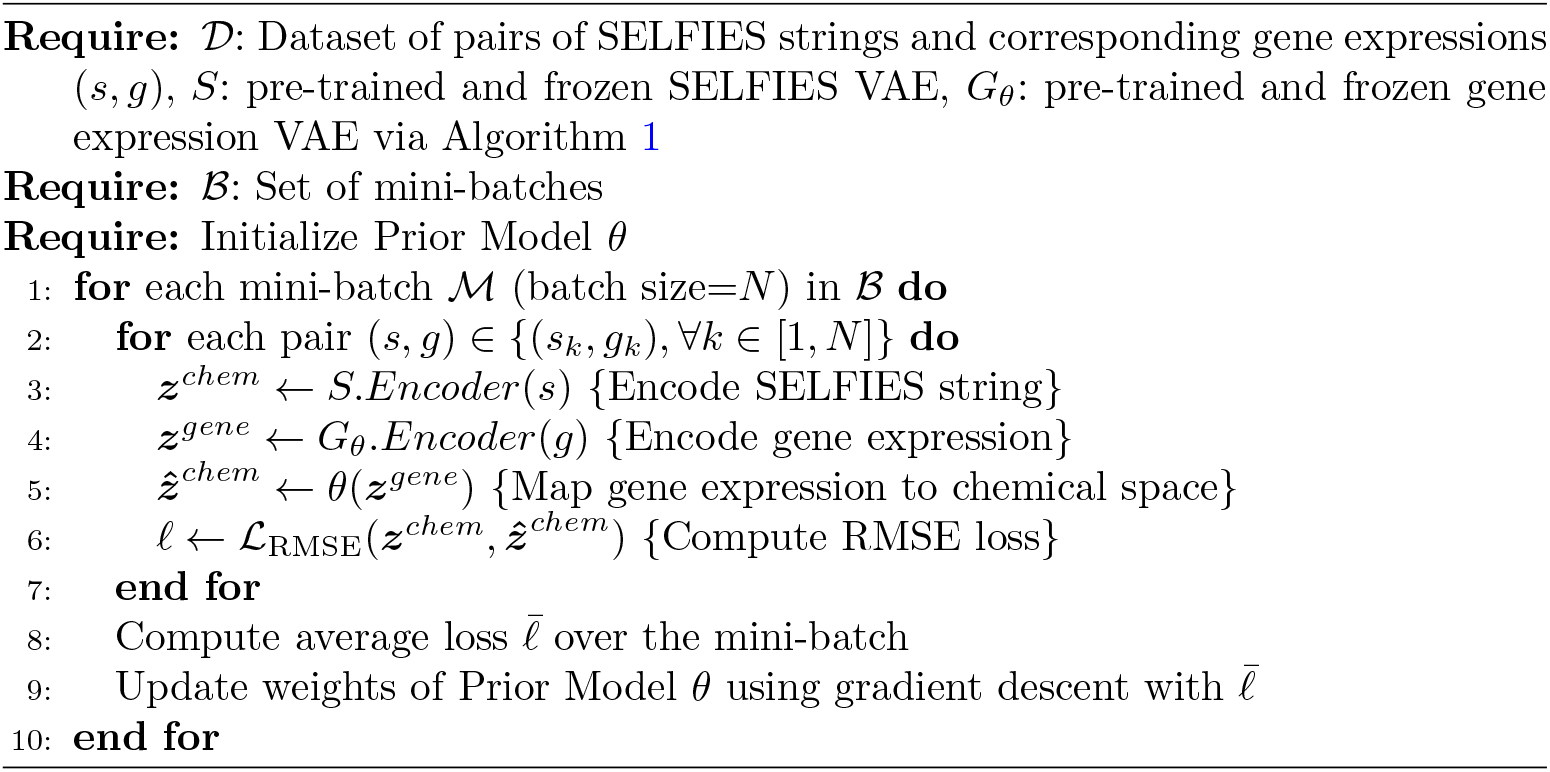

### Algorithm 3

Inference

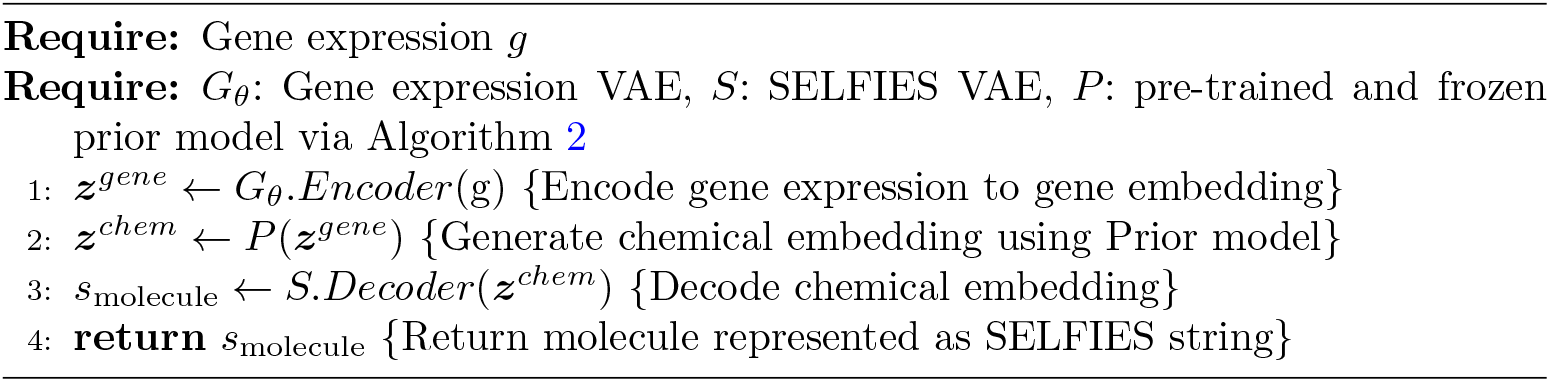

## Appendix

B Mean Gene Expression Signatures After Harmonizing Single Cell Dataset With L1000

**Fig. B1:**
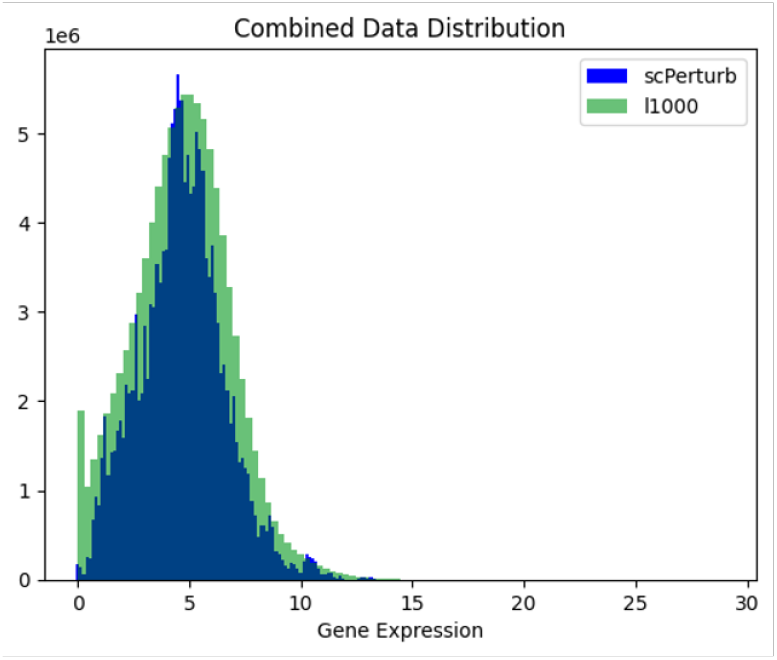
Mean gene expression signatures after harmonizing single cell dataset with L1000.

## Appendix

C Statistical Significance of Similarity Score Improvements

**Table C1:** P-values from one-sided Mann–Whitney U tests comparing MolGene-E against baseline models across different similarity metrics on the CRISPR perturbation dataset, where all comparisons are statistically significant (*p <* 0.05).

| Similarity | Baseline | P-value |
| --- | --- | --- |
| MACCS | MolGAN | $6.17 \times 10^{-4}$ |
| MACCS | GxVAE | $8.56 \times 10^{-7}$ |
| Fraggle | MolGAN | $2.30 \times 10^{-3}$ |
| Fraggle | GxVAE | $7.45 \times 10^{-3}$ |
| ErG | MolGAN | $5.18 \times 10^{-4}$ |
| ErG | GxVAE | $2.77 \times 10^{-4}$ |

**Table C2:** P-values from one-sided Mann–Whitney U tests comparing MolGene-E against baseline models across different similarity metrics on the Sciplex-3 dataset, where all comparisons are statistically significant (*p <* 0.05) except MolGene-E vs. MolGAN under ErG similarity.

| Similarity | Baseline | P-value |
| --- | --- | --- |
| MACCS | MolGAN | $3.98 \times 10^{-3}$ |
| MACCS | GxVAE | $8.04 \times 10^{-7}$ |
| Fraggle | MolGAN | $5.41 \times 10^{-4}$ |
| Fraggle | GxVAE | $1.98 \times 10^{-3}$ |
| ErG | MolGAN | $9.12 \times 10^{-2}$ |
| ErG | GxVAE | $1.74 \times 10^{-6}$ |

